# The longest intron rule

**DOI:** 10.1101/2023.10.02.560625

**Authors:** Pavel Dvorak, Viktor Hlavac, Vojtech Hanicinec, Bhavana Hemantha Rao, Pavel Soucek

## Abstract

Despite the fact that long introns mean an energy and time burden for eukaryotic cells, they play an irreplaceable role in the diversification and regulation of protein production. A general feature is the accumulation of the longest introns closer to the start of protein-coding genes. Our work aimed to more closely characterize the genes in which the longest intron is only located in the second or third third of the gene. Data on the lengths of all introns in genes were extracted from the genomes of 4 vertebrates (human, mouse, chicken and zebrafish), nematode worm and yeast. With the genes divided according to the relative position of the longest intron, gene set enrichment analyses were performed, the results of which were then confronted with the results of randomly generated gene sets of the same size. The analyses show that among the genes associated with biological processes of ion transport across membranes, cell signaling or the development of multicellular organisms, there is a greater representation of genes with the longest intron in the first third. Conversely, among the genes associated with the processes of RNA processing and ribosome biogenesis, there are more genes with the longest intron in the second or third third. It is a newly discovered characteristic with more general validity among multicellular organisms.

## Introduction

Although in the genomes of eukaryotic organisms we find a few percent of genes that consist of only one exon, the vast majority of genes are made up of both exons and introns. The length of individual exons and introns can vary by several orders of magnitude even within a single gene, and this variability primarily concerns the length of introns. The biological function of exons as gene segments encoding the sequence of amino acids in polypeptide chains, except for the less frequent cases of exons in the untranslated regions of genes, was recognized soon after they were defined. The functions of introns were revealed gradually, independently of the discussions about the time of their origin (introns-early versus introns-late theories), and it is still an incompletely explored issue [1]. Today, it is clear that introns play an irreplaceable role in increasing the diversity of proteins and in regulating the level of their gene expression [2,3].

The fact that the longest intron is significantly longer than the other introns in the gene and that it is often located closer to the 5’ end of the gene was already noted by the first works studying the structure of genes and genomes of sequenced eukaryotic organisms [4,5]. Information on the detailed distribution of intron lengths in the genes of various organisms has, however, been published so far only little and scattered. In our first study on this topic, we further defined the distribution of the longest introns in human genes - 64% of genes with the longest intron in the first third of all introns and 19 and 17% for the second and third third, with notable peaks for the middle and last introns of approx. 5 and 6%, respectively [6]. Moreover, we showed that localization of the longest intron in the second or third third is significantly more frequent for certain functionally related groups of genes, e.g. for DNA repair genes.

In this follow-up study, we extended the number of tested genes to the entire human genome and further tested five genomes of other organisms - mouse, chicken, zebrafish, nematode worm and yeast - with the intention of finding a possible more generally applicable rule of the relationship between the position of the longest intron and the biological function of the gene.

## Materials and methods

### Algorithm for calculating the lengths of all introns in protein-coding genes

In the first step, we obtained the positions of the beginnings (hereinafter referred to as Exon Start) and ends (Exon End) of all exons in the desired transcripts. For this, we used a query to the Ensembl database (https://www.ensembl.org/index.html; Release 108) [7] via the BioMart tool. In order to evaluate the most representative transcripts, we used MANE Select flags for Filters criteria in the human genome [8]. Considering that MANE Select is not available for other genomes, we sorted the transcripts in other organisms based on the selection of criteria Gene type - protein coding and at the same time Ensembl Canonical. In the event that even Ensembl Canonical flags were not available for the given genome, e.g. in Caenorhabditis elegans, we used APPRIS annotation instead. Among the data (Attributes) we queried for each transcript were Gene stable ID, Gene stable ID version, Transcript Stable ID, Gene name, Strand, Exon rank in transcript, Exon region start (bp, base pairs) and Exon region end (bp). In the second step, we used an in-house shell script to calculate the lengths of individual introns from the obtained data (the code is available in Supplementary Information S1). Genes were sorted not by their names (symbols) but by Gene or Transcript Stable IDs. We calculated intron lengths in bp for genes on the Forward strand according to the formula {[Exon(n+1)Start – Exon(n)End] – 1}; n are positive integers starting from 1. For genes on the Reverse strand, this formula was modified to {[Exon(n)Start – Exon(n+1)End] – 1}. The script than created a table with the lengths of all introns for each protein-coding gene and searched for the position of the longest intron in the gene. In particular, we used AWK language to perform the following steps: 1) Calculate the lengths of the introns; 2) Extract Gene names if they are available; 3) Indicate the longest intron; 4) Calculate the relative position of the longest intron (the ratio of the position of the longest intron to the total number of introns in the given gene) and Bourne Again Shell (BASH) to create a matrix of Gene or Transcript Stable IDs versus the lengths of the introns in bp. The script can handle various delimiters and require an input file exported from Ensembl BioMart query with either Gene Stable ID Version or Transcript stable ID as the main unique identifier. The first seven columns in the file must be: (1) Gene stable ID, (2) Gene or Transcript stable ID (version), (3) Gene name, (4) Strand, (5) Exon rank in transcript, (6) Exon region start and (7) Exon region end. The primary data thus obtained, which we analyzed in this study, for individual organisms – Homo sapiens (human), Mus musculus (mouse), Gallus gallus (chicken), Danio rerio (zebrafish), Caenorhabditis elegans (worm) and Saccharomyces cerevisiae (yeast) – are stored in Supplementary Information Tables S2-7, respectively.

### Gene set enrichment analysis (GSEA)

For GSEA, we selected genes that contain at least three introns from each studied genome, with the exception of yeast’s genome, and further divided these genes into three subgroups according to the relative position of the longest intron (defined above). The position of the longest intron in the first third of all introns means that this ratio is in the interval (0;0.33]. Similarly, for the second third it is in the interval (0.33;0.66] and for the third third in (0.66;1]. In yeast, where there is only a minimal number of genes with more than one intron, we divided all protein-coding genes into only two tested groups: 1) without introns and 2) with introns. Our input (primary) data for GSEA can be obtained from Table S8.

We performed GSEA in parallel using two web platforms – g:Profiler (https://biit.cs.ut.ee/gprofiler/gost) [9] and ShinyGO (http://bioinformatics.sdstate.edu/go/) [10], taking into account the procedure recommended in the work of Reimand et al. [11]. With the g:Profiler program, we used the option to analyze multiple gene files simultaneously (*Run as multiquery* option), other possible options for setting the result parameters were left in the default settings, even with the ShinyGO program. We used the AmiGO 2 project (http://amigo.geneontology.org/amigo/landing) [12] for better orientation in the hierarchical structure of GO terms (Gene Ontology; http://geneontology.org/) [13] in the resulting lists of terms and the tool for creating Venn diagrams Multiple List Comparator (https://molbiotools.com/listcompare.php) [14] for finding common terms between these lists.

KEGG diagrams (Kyoto Encyclopedia of Genes and Genomes; https://www.genome.jp/kegg/) [15] were created with the help of Pathview (https://pathview.uncc.edu/) [16].

### Random gene lists

In order to obtain comparable control data, we generated random sets of genes using the RSAT web server (http://rsat.sb-roscoff.fr/index.php) [17] and performed the same analyses as the primary data. For each of the multicellular organisms tested, we generated three sets, for yeast only two, always with the same number of genes as for adequate primary data. Control data is stored in Table S9.

## Results

### Characteristics of the studied genomes

We analyzed 4 genomes of representatives of the vertebrate subphylum (human, mouse, chicken and zebrafish) and one genome each from the phylum Nematoda (worm) and division Ascomycota (yeast). Of the mentioned vertebrates, the chicken has the fewest protein-coding genes (approx. 17,000) and the zebrafish the most (30,000), the human and the mouse have 19 and 22 thousand genes respectively. The median number of introns in a gene is similar among all four vertebrates, i.e. 6 or 7, and approximately 80% of genes carry 3 or more introns. Human and mouse have approximately 60% of genes that have their longest introns located in the first third and 20% each of genes that have their longest introns located in the second and third thirds. In chicken and zebrafish, this ratio is skewed towards a higher percentage of genes that have the longest intron in the third third, i.e. 24 and 32%, respectively. In the nematode worm, the number of protein-coding genes is also about 20 thousand, but the median number of introns in a gene is only 4, and the number of genes with the longest intron in the first third is even lower (about 40%). From the point of view of introns, the unicellular yeast genome, with approximately 7,000 genes, is fundamentally different.

96% of the genes do not have a single intron present, and the remaining 4% are genes with one intron (only 13 yeast genes carry two introns and 5 genes more than two). Therefore, the division into groups according to the position of the longest intron is not relevant here. More precise values are shown in Table 1.

**Table 1:**
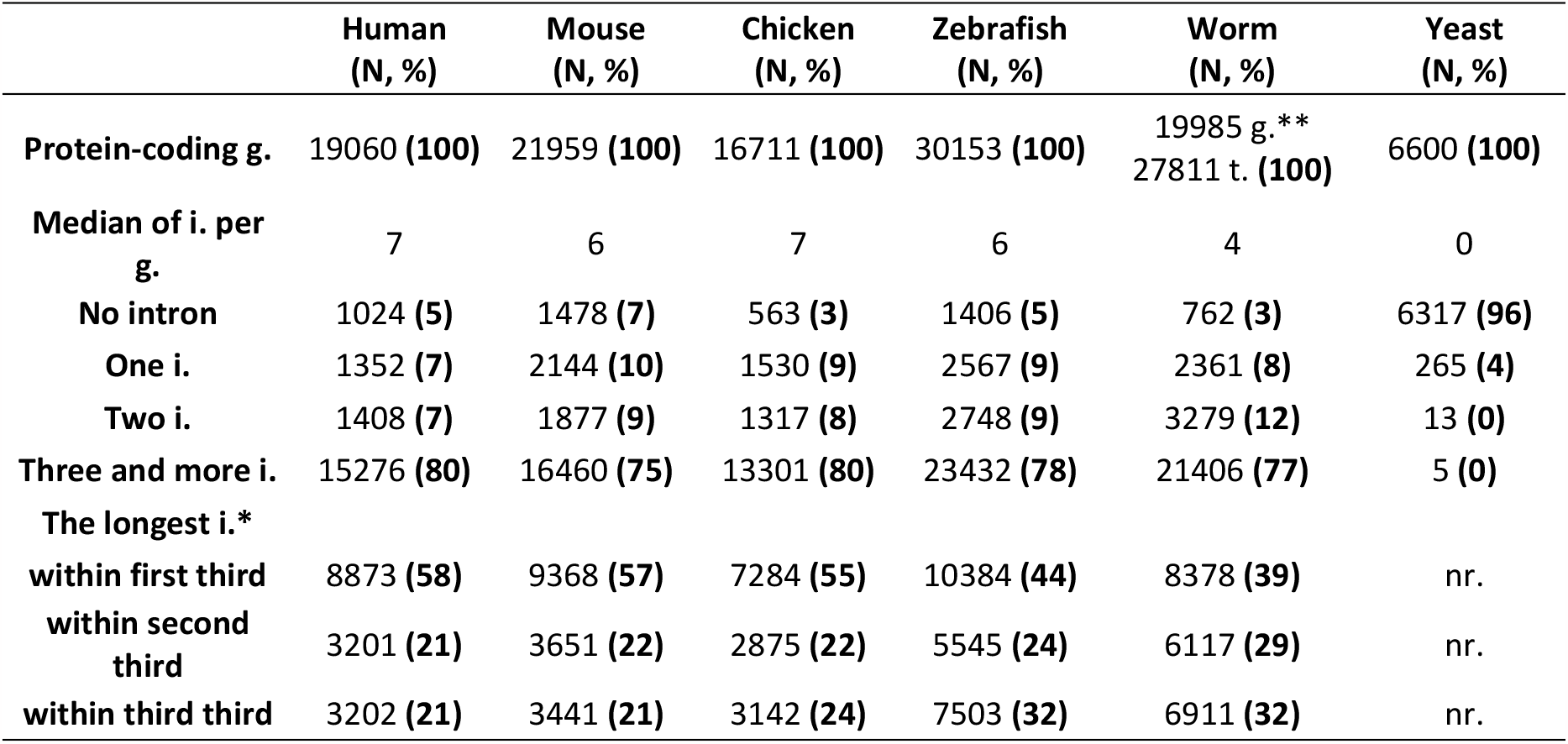
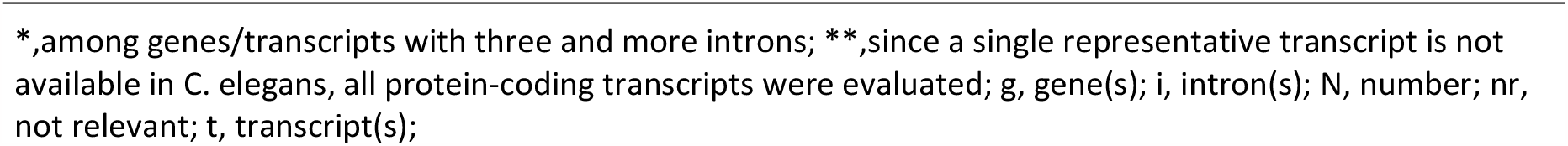
Numbers and localization of introns within protein-coding genes of six studied eukaryotic genomes.

|  | Human<br>(N, %) | Mouse<br>(N, %) | Chicken<br>(N, %) | Zebrafish<br>(N, %) | Worm<br>(N, %) | Yeast<br>(N, %) |
| --- | --- | --- | --- | --- | --- | --- |
| **Protein-coding g.** | 19060 (100) | 21959 (100) | 16711 (100) | 30153 (100) | 19985 g. **<br>27811 t. (100) | 6600 (100) |
| **Median of i. per**<br>**g.** | 7 | 6 | 7 | 6 | 4 | 0 |
| **No intron** | 1024 (5) | 1478 (7) | 563 (3) | 1406 (5) | 762 (3) | 6317 (96) |
| **One i.** | 1352 (7) | 2144 (10) | 1530 (9) | 2567 (9) | 2361 (8) | 265 (4) |
| **Two i.** | 1408 (7) | 1877 (9) | 1317 (8) | 2748 (9) | 3279 (12) | 13 (0) |
| **Three and more i.** | 15276 (80) | 16460 (75) | 13301 (80) | 23432 (78) | 21406 (77) | 5 (0) |
| **The longest i.\*** |  |  |  |  |  |  |
| **within first third** | 8873 (58) | 9368 (57) | 7284 (55) | 10384 (44) | 8378 (39) | nr. |
| **within second**<br>**third** | 3201 (21) | 3651 (22) | 2875 (22) | 5545 (24) | 6117 (29) | nr. |
| **within third third** | 3202 (21) | 3441 (21) | 3142 (24) | 7503 (32) | 6911 (32) | nr. |

\*,among genes/transcripts with three and more introns; \*\*,since a single representative transcript is not available in *C. elegans*, all protein-coding transcripts were evaluated; g, gene(s); i, intron(s); N, number; nr, not relevant; t, transcript(s);

### Overrepresentation of specific Gene Ontology terms among genes that have the longest intron localized in the first part of the gene

The analysis focused on GO terms from the Biological Process category characterizing genes that have the longest intron located in the first third of the gene found a large number of terms exclusively overrepresented in this subgroup, especially in the human and mouse genomes (Table S10). The intersection of all these terms among the five studied multicellular organisms contained the following 12 terms: (1) *cell morphogenesis*, (2) *dephosphorylation*, (3) *inorganic cation transmembrane transport*, (4) *inorganic ion transmembrane transport*, (5) *metal ion transport*, (6) *monoatomic cation transmembrane transport*, (7) *monoatomic cation transport*, (8) *monoatomic ion transmembrane transport*, (9) *monoatomic ion transport*, (10) *multicellular organismal process*, (11) *sodium ion transport* and (12) *transmembrane transport*. The term *transmembrane transport* (12) is superior to the terms (3), (4), (6) and (8), and the term *monoatomic ion transport* (9) is superior to the terms (5), (6), (7), (8) and (11); the other terms are not closely related. Another 121 terms (shown in Table S11) were common to four of the tested multicellular organisms. P values for selected terms are demonstrated in Table 2.

**Table 2:**
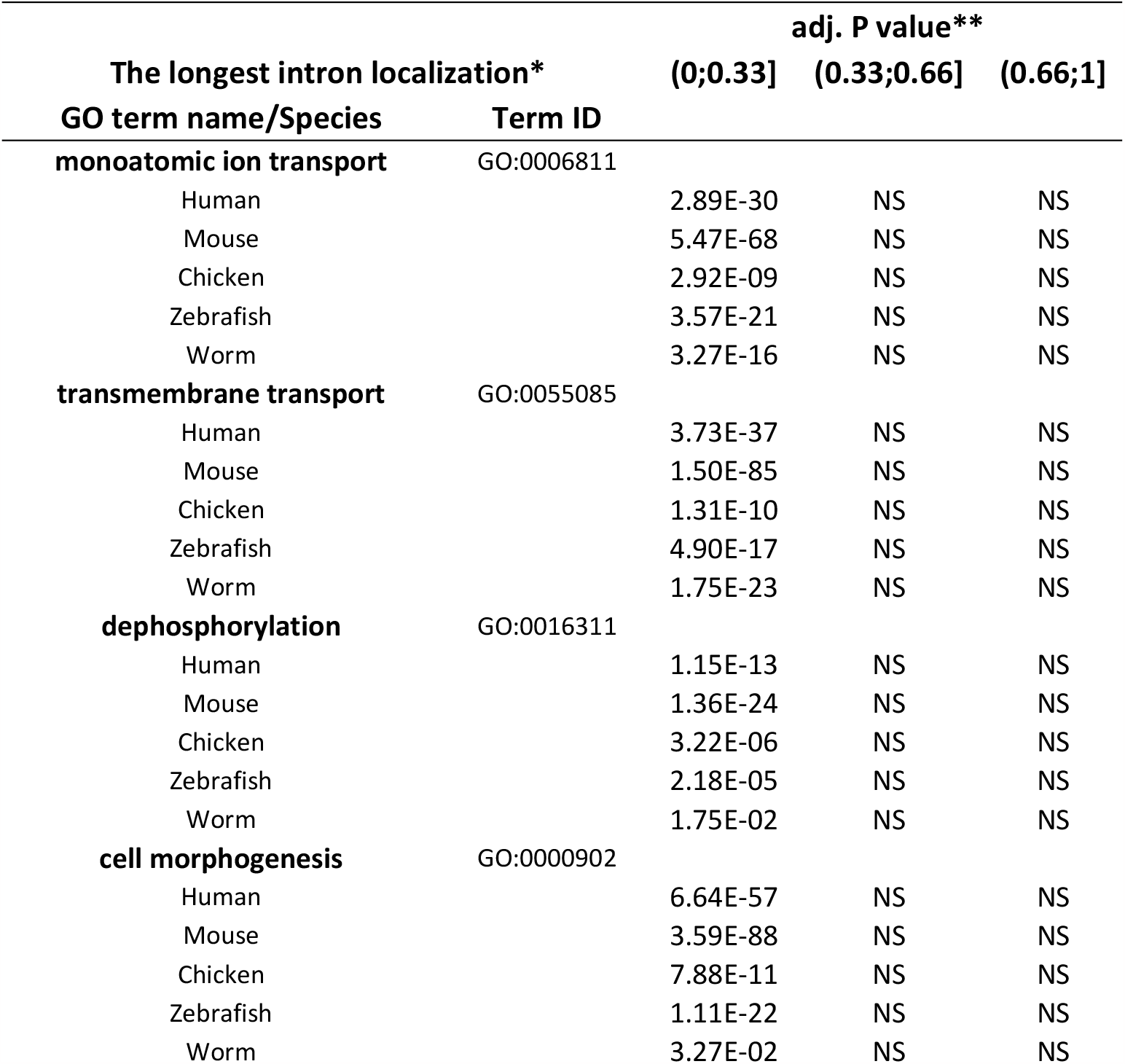
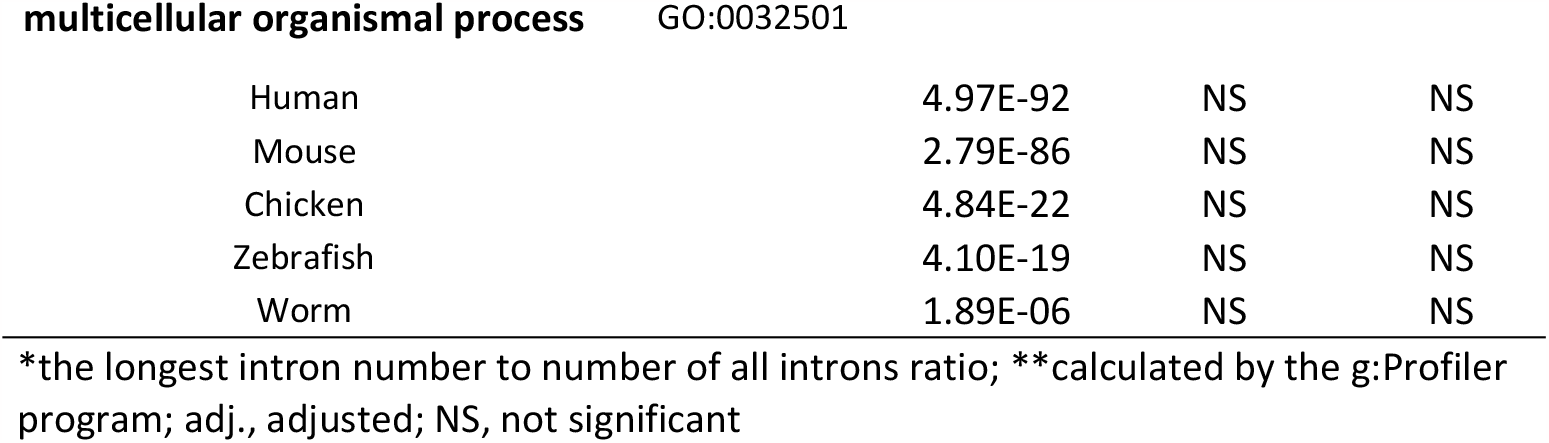
Selected Gene Ontology (GO) terms exclusively overrepresented in genes with the longest intron localized in the first part of the gene.

### Overrepresentation of specific Gene Ontology terms among genes that have the longest intron localized in the second or third part of the gene

In a similar manner, in all evaluated multicellular organisms, three GO terms from the Biological Process category – (1) *ncRNA metabolic process*, (2) *ncRNA processing* and (3) *ribosome biogenesis* - showed a significantly higher occurrence exclusively among genes in which the longest intron is located in the second or third part. Even in the case of yeast, these three GO terms are overrepresented among genes with introns compared to genes without introns (Table 3). The term No. (1) is superior to (2). A number of other terms showed the same characteristics in several of the studied genomes. In particular, the following five terms - *RNA modification, chromosome organization, rRNA metabolic process, tRNA metabolic process* and *translation* - were common to the four multicellular organisms and another twelve terms including *DNA repair, cilium organization, methylation, nuclear division* and *rRNA processing* to three organisms (Table S12). All these results for the longest introns localized in the second or third part are listed in Table S13 including P values of individual GO terms.

**Table 3:**
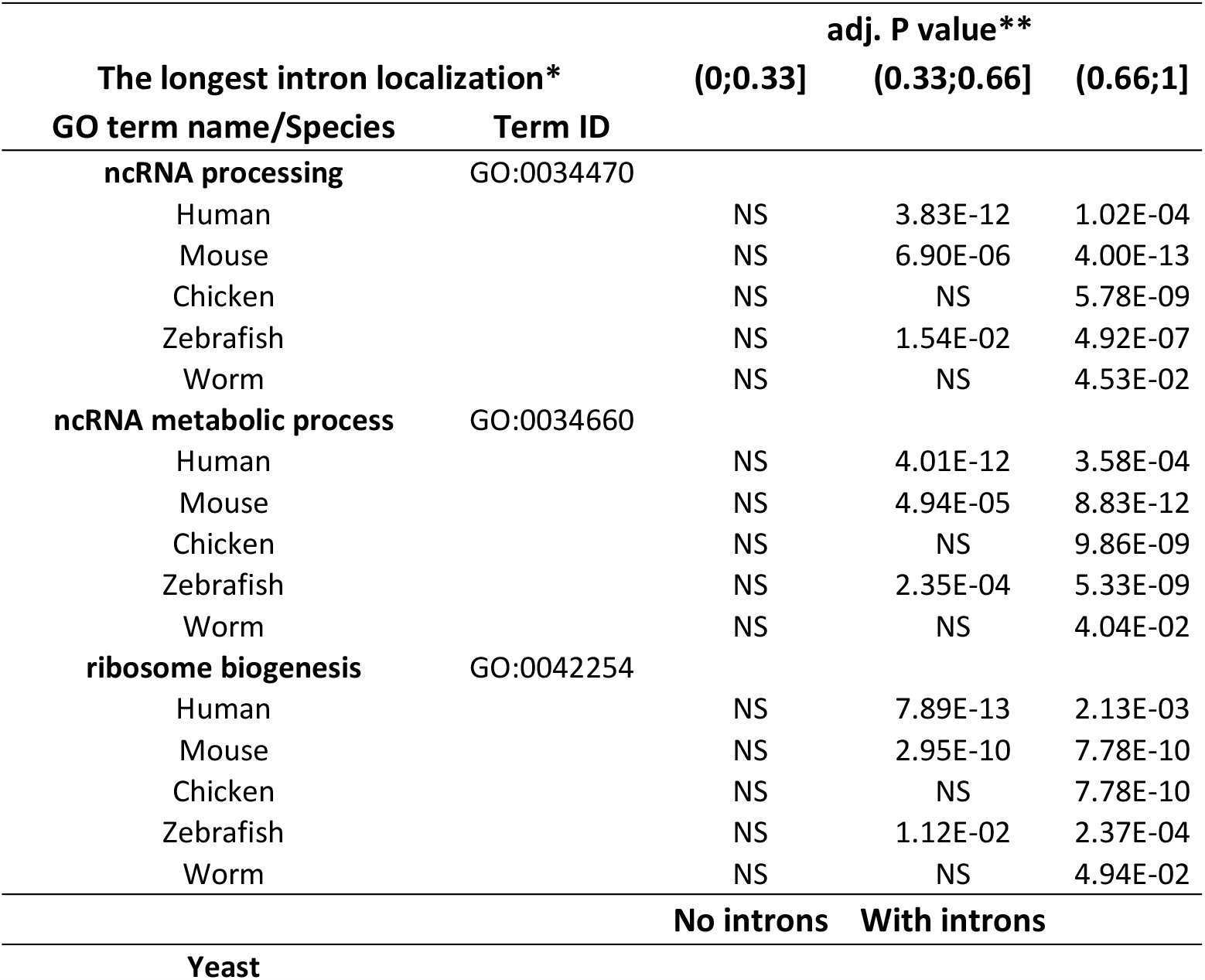
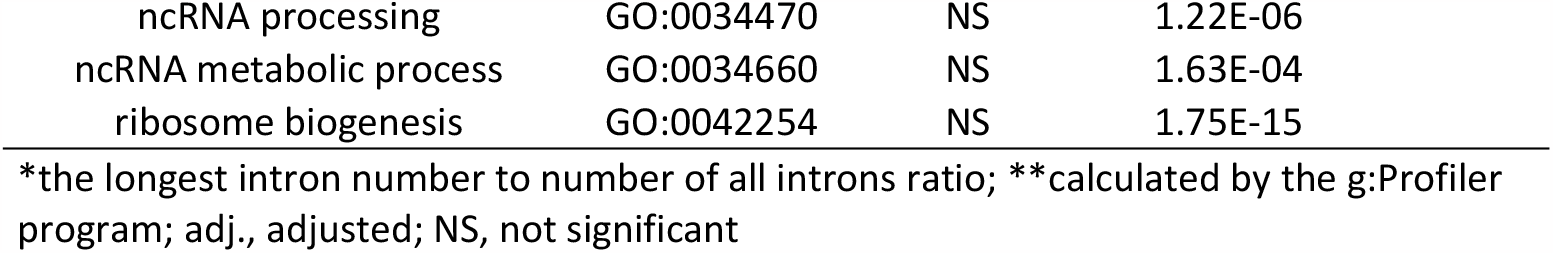
Selected Gene Ontology (GO) terms exclusively overrepresented in genes with the longest intron localized in the second or third part of the gene.

|  |  | adj. P value** |  |  |
| --- | --- | --- | --- | --- |
| The longest intron localization* |  | (0;0.33] | (0.33;0.66] | (0.66;1] |
| GO term name/Species | Term ID |  |  |  |
| **ncRNA processing** | GO:0034470 |  |  |  |
| Human |  | NS | 3.83E-12 | 1.02E-04 |
| Mouse |  | NS | 6.90E-06 | 4.00E-13 |
| Chicken |  | NS | NS | 5.78E-09 |
| Zebrafish |  | NS | 1.54E-02 | 4.92E-07 |
| Worm |  | NS | NS | 4.53E-02 |
| **ncRNA metabolic process** | GO:0034660 |  |  |  |
| Human |  | NS | 4.01E-12 | 3.58E-04 |
| Mouse |  | NS | 4.94E-05 | 8.83E-12 |
| Chicken |  | NS | NS | 9.86E-09 |
| Zebrafish |  | NS | 2.35E-04 | 5.33E-09 |
| Worm |  | NS | NS | 4.04E-02 |
| **ribosome biogenesis** | GO:0042254 |  |  |  |
| Human |  | NS | 7.89E-13 | 2.13E-03 |
| Mouse |  | NS | 2.95E-10 | 7.78E-10 |
| Chicken |  | NS | NS | 7.78E-10 |
| Zebrafish |  | NS | 1.12E-02 | 2.37E-04 |
| Worm |  | NS | NS | 4.94E-02 |
|  |  | **No introns** | **With introns** |  |
| **Yeast** |  |  |  |  |

|  |  |  |  |
| --- | --- | --- | --- |
| ncRNA processing | GO:0034470 | NS | 1.22E-06 |
| ncRNA metabolic process | GO:0034660 | NS | 1.63E-04 |
| ribosome biogenesis | GO:0042254 | NS | 1.75E-15 |

\*the longest intron number to number of all introns ratio; \*\*calculated by the g:Profiler program; adj., adjusted; NS, not significant

### Increased occurrence of genes included in certain KEGG pathways depending on the position of the longest intron in the gene

The analysis focused on the participation of genes, divided according to the position of the longest intron, in specific KEGG pathways brought additional results. The number of genes with the longest intron in the first third was significantly increased, in all 5 tested multicellular organisms, for the following 3 pathways: Calcium signaling pathway, FoxO signaling pathway and Purine metabolism. Additional thirty-eight pathways, including Apoptosis, Adherens, Tight and Gap junctions, ErbB, Hedgehog and Insulin signaling pathways, and Carbon and Glycerolipid metabolisms, met statistical significance in 4 organisms (Table S14). Conversely, for genes with the longest intron in the second or third third, the most represented pathways were: Biosynthesis of cofactors, Ribosome, Spliceosome, Cell cycle, Proteasome and Ribosome biogenesis in eukaryotes (Table S15). A demonstrative selection of KEGG pathways is shown in Table 4, and more detailed results can be found in Tables S16 and S17. Genes from the Spliceosome pathway, which have the longest intron located in the second or third third of the gene in the model organism zebrafish, are highlighted in Figure 1.

**Table 4:** Selected KEGG pathways overrepresented among genes divided according to the position of the longest intron.

| KEGG pathway | adj. P value* |  |  |  |  |
| --- | --- | --- | --- | --- | --- |
|  | Human | Mouse | Chicken | Zebrafish | Worm |
| **The longest intron within first third** |  |  |  |  |  |
| Calcium signaling pathway | 4.9E-10 | 2.3E-05 | 1.3E-07 | 6.1E-13 | 3.5E-05 |
| FoxO signaling pathway | 1.3E-04 | 1.9E-02 | 3.5E-04 | 2.6E-05 | 1.6E-02 |
| ErbB signaling pathway | 2.9E-10 | 7.5E-08 | 9.7E-05 | 5.6E-15 | NS |
| Purine metabolism | 2.0E-07 | 9.8E-06 | 1.0E-02 | 1.7E-10 | 9.1E-04 |
| Carbon metabolism | 4.0E-07 | 1.6E-04 | 2.9E-02 | 4.6E-02 | NS |
| Adherens junction | 1.1E-11 | 8.2E-09 | 4.4E-09 | 1.1E-16 | NS |
| Gap junction | 4.4E-08 | 2.8E-08 | 4.4E-03 | 1.9E-13 | NS |
| Apoptosis | 1.3E-06 | 5.0E-05 | 3.7E-02 | 2.7E-06 | NS |
| **The longest intron within second or third third** |  |  |  |  |  |
| Biosynthesis of cofactors | 1.9E-02 | 1.3E-02 | 4.2E-04 | 7.8E-04 | 4.0E-02 |
| Ribosome | 2.4E-17 | 1.2E-11 | 6.9E-09 | 1.4E-11 | NS |
| Spliceosome | 7.5E-04 | 3.9E-02 | 1.2E-07 | 6.7E-10 | 3.5E-02 |
| Cell cycle | 1.6E-04 | 2.0E-02 | 4.2E-04 | 1.1E-07 | NS |
| Proteasome | 4.3E-02 | 8.5E-03 | 1.5E-02 | 5.7E-06 | NS |
| Ribosome biogenesis in eukaryotes | 2.9E-02 | NS | 1.6E-05 | 1.1E-04 | 2.9E-02 |

\*calculated by the ShinyGO program; adj., adjusted; NS, not significant

**Figure 1:**
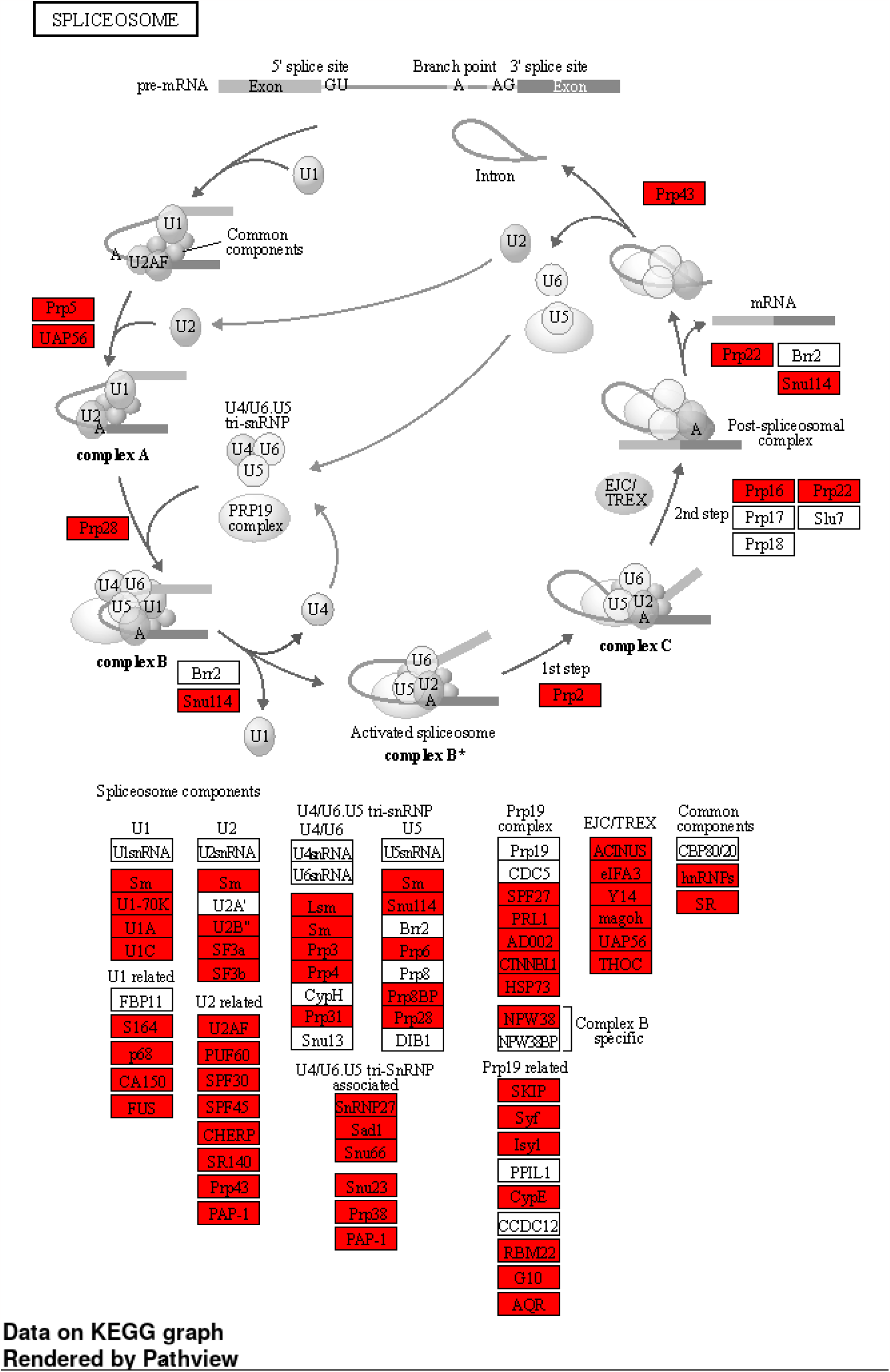
The Spliceosome pathway in the model organism zebrafish. The genes, which have the longest intron located in the second or third third of the gene are highlighted in red.

### Results from control datasets

No KEGG pathway was significantly increased in the control data, nor were any of the GO terms belonging to the Biological Process category, which were overrepresented in the primary data. Only a few terms from other categories (e.g. cytoplasm and protein binding) met the criteria for statistical significance in random gene control sets, but only in single cases without repetition in another organism.

## Discussion

Although longer introns represent an energy and time burden for the genetic apparatus of the cell trying to efficiently transfer information from its storage to concrete implementation, they perform important and apparently irreplaceable tasks in this process. Our presented analyses show that a characteristic position of the longest intron can be found in genes with certain biological functions. The results in five multicellular and one unicellular eukaryotic organisms suggest the more general validity of this previously unreported characteristic.

We have evidence that longer introns contain higher densities of conserved sites [18] and various regulatory elements [19,20]. Also, the presence of longer introns closer to the 5’ end of eukaryotic genes is a well-known phenomenon [21], which is explained by a greater occurrence of regulatory elements and thus a greater restriction of selectivity in these introns.

The current work, on the other hand, aimed to focus on genes that have the longest intron located in the second or third third of the gene. We specified the proportion of these protein-coding genes in the genomes of five multicellular organisms. Such genes make up 61% of the nematode worm genome, 56% in zebrafish, 45% in chicken, and only around 40% in humans and mice. It appears that the proportion of genes with the longest intron in the first third of the gene increases significantly with the complexity of multicellular organisms.

It is evident from our results that among the genes associated with the processes of cell signaling, export from cell, cell junction organization, peptidyl-amino acid modification or developmental growth, there is a greater representation of genes with the longest intron in the first third. And this characteristic is again more evident in more complex organisms. Conversely, among the genes associated with nuclear division, ribosome biogenesis, macromolecule methylation or cytoplasmic translation, there are more genes with the longest intron in the second or third third; with more general validity among studied organisms.

The term ncRNA (non-coding RNA) is superior both to RNA molecules involved in the formation of proteins according to the information stored in mRNA (rRNA, tRNA, snRNA and snoRNA), and to RNA molecules with various other regulatory functions (snoRNA, miRNA, siRNA and others), see e.g., the work of Mattick and Makunin [22]. Thus, the term ncRNA processing encompasses a whole constellation of more broadly related events. However, the core RNA processing mechanisms and machinery are highly conserved, with dysregulation of constitutive patterns often having severe cellular consequences [23]. Some authors even consider RNA processing as a relic that can be traced back either to the RNA world that preceded both DNA and encoded protein synthesis or to the later ribonucleoprotein (RNP) world [24]. The differences in the length structure of the introns, discussed by us, in these genes are most likely related to this high evolutionary conservation.

The influence of introns on the regulation of genes for ribosomal proteins and the fact that these genes carry introns even in the otherwise intron-poor yeast genome have already been mentioned in the published literature [25]. The presence of introns substantially increases the complexity of ribosomal protein gene expression as they variably slow the expression cycle, and in addition, many introns can contain non-coding RNA involved in other layers of regulation [2]. In current works on ribosomal proteins, their other functions beyond the framework of ribosome formation and maintenance are emphasized, e.g. activation of p53-dependent pathway in response to stress [26,27]. Thus, the significance of the longest intron feature of these genes highlighted by us may also turn out to be broader and awaits further investigation.

## Conclusions

In eukaryotic organisms, genes related to more complex functions of multicellular organisms tend to locate the longest intron in the first third of the gene, while genes related to very ancient and evolutionarily conserved functions related to RNA processing show the position of the longest intron mainly in the second or third third.

## Availability of data and materials

The datasets - code as well as all Supplementary Tables - generated during and/or analysed during the current study are available in the Zenodo repository, DOI: 10.5281/zenodo.8398541, https://zenodo.org/record/8398541.

## Declarations

### Authors’ contributions

**Pavel Dvorak:** Conceptualization, Methodology, Validation, Formal analysis, Writing - Original Draft, Visualization, Funding acquisition; **Viktor Hlavac:** Methodology, Programming, Data Analysis, Writing - Original Draft; **Vojtech Hanicinec:** Software, Formal analysis, Data Curation, Writing - Original Draft; **Bhavana Hemantha Rao:** Software, Formal analysis, Data Curation, Writing - Original Draft; **Pavel Soucek:** Writing - Review & Editing, Supervision, Funding acquisition;

## References

1. Rogozin, I. B., Carmel, L., Csuros, M. & Koonin, E. V. Origin and evolution of spliceosomal introns. Biol Direct 7, 11 (2012).

2. Chorev, M. & Carmel, L. The Function of Introns. Front. Gene. 3, (2012).

3. Parenteau, J. & Abou Elela, S. Introns: Good Day Junk Is Bad Day Treasure. Trends in Genetics 35, 923–934 (2019).

4. Hawkin, J. D. A survey on intron and exon lengths. Nucl Acids Res 16, 9893–9908 (1988).

5. Sakurai, A. et al. On biased distribution of introns in various eukaryotes. Gene 300, 89–95 (2002).

6. Dvorak, P., Hanicinec, V. & Soucek, P. The position of the longest intron is related to biological functions in some human genes. Front. Genet. 13, 1085139 (2023).

7. Cunningham, F. et al. Ensembl 2022. Nucleic Acids Research 50, D988–D995 (2022).

8. Morales, J. et al. A joint NCBI and EMBL-EBI transcript set for clinical genomics and research. Nature 604, 310–315 (2022).

9. Raudvere, U. et al. g:Profiler: a web server for functional enrichment analysis and conversions of gene lists (2019 update). Nucleic Acids Research 47, W191–W198 (2019).

10. Ge, S. X., Jung, D. & Yao, R. ShinyGO: a graphical gene-set enrichment tool for animals and plants. Bioinformatics 36, 2628–2629 (2020).

11. Reimand, J. et al. Pathway enrichment analysis and visualization of omics data using g:Profiler, GSEA, Cytoscape and EnrichmentMap. Nat Protoc 14, 482–517 (2019).

12. Carbon, S. et al. AmiGO: online access to ontology and annotation data. Bioinformatics 25, 288–289 (2009).

13. Ashburner, M. et al. Gene Ontology: tool for the unification of biology. Nat Genet 25, 25–29 (2000).

14. Jia, A., Xu, L. & Wang, Y. Venn diagrams in bioinformatics. Briefings in Bioinformatics 22, bbab108 (2021).

15. Kanehisa, M., Furumichi, M., Sato, Y., Ishiguro-Watanabe, M. & Tanabe, M. KEGG: integrating viruses and cellular organisms. Nucleic Acids Research 49, D545–D551 (2021).

16. Luo, W., Pant, G., Bhavnasi, Y. K., Blanchard, S. G. & Brouwer, C. Pathview Web: user friendly pathway visualization and data integration. Nucleic Acids Research 45, W501–W508 (2017).

17. Santana-Garcia, W. et al. RSAT 2022: regulatory sequence analysis tools. Nucleic Acids Research 50, W670–W676 (2022).

18. Shin, S.-H. & Choi, S. S. Lengths of coding and noncoding regions of a gene correlate with gene essentiality and rates of evolution. Genes Genom 37, 365–374 (2015).

19. Majewski, J. & Ott, J. Distribution and Characterization of Regulatory Elements in the Human Genome. Genome Res. 12, 1827–1836 (2002).

20. Park, S. G., Hannenhalli, S. & Choi, S. S. Conservation in first introns is positively associated with the number of exons within genes and the presence of regulatory epigenetic signals. BMC Genomics 15, 526 (2014).

21. Bradnam, K. R. & Korf, I. Longer First Introns Are a General Property of Eukaryotic Gene Structure. PLoS ONE 3, e3093 (2008).

22. Mattick, J. S. & Makunin, I. V. Non-coding RNA. Human Molecular Genetics 15, R17–R29 (2006).

23. Pai, A. A. & Luca, F. Environmental influences on RNA processing: Biochemical, molecular and genetic regulators of cellular response. WIREs RNA 10, (2019).

24. Penny, D., Hoeppner, M. P., Poole, A. M. & Jeffares, D. C. An Overview of the Introns-First Theory. J Mol Evol 69, 527–540 (2009).

25. Petibon, C., Malik Ghulam, M., Catala, M. & Abou Elela, S. Regulation of ribosomal protein genes: An ordered anarchy. WIREs RNA 12, (2021).

26. Jiao, L. et al. Ribosome biogenesis in disease: new players and therapeutic targets. Sig Transduct Target Ther 8, 15 (2023).

27. Kang, J. et al. Ribosomal proteins and human diseases: molecular mechanisms and targeted therapy. Sig Transduct Target Ther 6, 323 (2021).

